# Exploration of Chemical Space with Partial Labeled Noisy Student Self-Training for Improving Deep Learning Performance: Application to Drug Metabolism

**DOI:** 10.1101/2020.08.06.239988

**Authors:** Yang Liu, Hansaim Lim, Lei Xie

## Abstract

**Motivation:** Drug discovery is time-consuming and costly. Machine learning, especially deep learning, shows a great potential in accelerating the drug discovery process and reducing its cost. A big challenge in developing robust and generalizable deep learning models for drug design is the lack of a large amount of data with high quality and balanced labels. To address this challenge, we developed a self-training method PLANS that exploits millions of unlabeled chemical compounds as well as partially labeled pharmacological data to improve the performance of neural network models.

**Result:** We evaluated the self-training with PLANS for Cytochrome P450 binding activity prediction task, and proved that our method could significantly improve the performance of the neural network model with a large margin. Compared with the baseline deep neural network model, the PLANS-trained neural network model improved accuracy, precision, recall, and F1 score by 13.4%, 12.5%, 8.3%, and 10.3%, respectively. The self-training with PLANS is model agnostic, and can be applied to any deep learning architectures. Thus, PLANS provides a general solution to utilize unlabeled and partially labeled data to improve the predictive modeling for drug discovery.

**Availability:** The code that implements PLANS is available at https://github.com/XieResearchGroup/PLANS

## 1. Introduction

Recent advances in deep learning have shown promise in accelerating drug discovery (Chen *et al*., 2018). However, several challenges remain in the successful application of deep learning to drug discovery. A general problem for almost all supervised deep learning is that the amount of high-quality labeled data is usually limited. To train an accurate, robust, and generalizable model using the deep learning, it is not surprising to use millions even billions of labeled training data. Even the most advanced high throughput experimental methods today cannot meet the huge data demand. Thus, methods are needed to sufficiently exploit the labeled pharmacological data and unlabeled chemical space to improve the performance of deep learning models. Autoencoder (AE) (Rumelhart *et al*., 1986; Kipf and Welling, 2016a) and variant autoencoder (VAE) (Kingma and Welling; Kipf and Welling, 2016b) are widely used models which, after training with unlabeled data, can better represent input data features by mapping the sparse sample space to a continuous latent space. Multi-task models try to find a better latent space by combining datasets with labels for different tasks (Collobert and Weston, 2008; Ramsundar *et al*., 2015). Self-training method, on the other hand, use the “teacher” model trained with small amount data to label the unlabeled data and recursively train “student” models that have better properties than the “teacher” model (Hinton *et al*., 2015). Recently, Google developed a “Noisy Student” self-training method, in which they used a simple teacher model to train a series of student models with gradually increasing complexity (Xie *et al*.). In their experiment, the Noisy Student trained models showed not only better performance, but also more robust. In spite of their successes in imaging processing and Natural Language Processing, few works have applied the concept of self-training to chemical compound screening and other machine learning tasks in drug discovery given that billions of chemical compounds are not associated with any labels. In addition to huge amount of unlabeled data, bioassay data is often highly imbalanced. Conventional over-sampling or over-sampling techniques such as SMOTE (Chawla *et al*., 2011), which rely on the similarity between samples, maybe not sufficient in addressing this problem. On one hand, a small modification in a chemical structure could result in a dramatic change in bioactivity (Stumpfe and Bajorath, 2012). On the other hand, highly dissimilar chemical compounds could have similar bioactivity (Böhm *et al*., 2004). Therefore, new techniques are needed to address imbalanced sample problem.

In this work, for the first time, we design a self-training strategy to explore unlabeled chemical space for improving predictive modeling in drug discovery. We develop a new self-training method: Partially LAbeled Noisy Student (PLANS). Moreover, we use the self-training with PLANS to address the problem of sample imbalance. To demonstrate the value of PLANS, we apply our method to the prediction of Cytochrome P450 (CYP450) chemical binding profile. CYP450 is a protein enzyme family catalyze hydroxyl group incorporation with heme as a cofactor. In human, CYP450s, as terminal oxidase in electron transferring chain, participate in a wide range of metabolism processes including hormone synthesis, fatty acids synthesis, steroids oxidization, etc. (McDonnell and Dang, 2013). CYP450s play key roles in drug metabolism. The activation or deactivation of approximate 75 % drugs is mediated by CYP450s (Guengerich, 2008). The human genome has 57 CYP450 genes expressing proteins share similar folding (Nebert *et al*., 2013). Therefore, CYP450s are one of the major reasons of adverse drug interactions. Furthermore, the polymorphism of CYP450s accounts for the individual difference in drug responses. Study in the binding profile of drugs to CYP450s is also a fruitful source for pharmacogenomics. The binding profile of a large number of chemicals with CYP450 remains unknown. However, few machine learning algorithms are able to reliably predict CYP450-drug interactions. Five CYP450s, CYP1A2, CYP2C9, CYP2C19, CYP2D6, and CYP3A4 are chosen to be the targets for chemical compounds in this work, which play the most important roles in drug metabolism (Cupp and Tracy, 1998). Comprehensive benchmark studies demonstrate that the self-training with PLANS significantly improve the performance of the CYP450 binding profile prediction. PLANS is model agnostic. They could be applied to other deep learning tasks for drug design using any neural network architectures and with limited amount of noisy data.

## 2. Materials and methods

### 2.1. Overview of PLANS

As shown in Figure 1, the work flow of PLANS includes teacher initialization, iterative training, and final testing. In the initialization stage, we train the first teacher model with only the fully labeled data. Then, in the iterative training stage, we use the trained teacher model to generate labels for partially labeled and unlabeled datasets. Data balancing can be introduced at this stage if needed. The fully labeled data and data with inferred labels are combined to train a noisy student model. Then the noisy student model is used as a new teacher model to generate labels for the partially labeled and unlabeled data. This step can be repeated until the performance of the noisy student model does not further improve. In the testing stage, we simply use the last student model to predict labels for the testing data. In the whole process, the noises are introduced by label mixup (see below) and dropout, and used only when training the student models. They are disabled when using the model as teacher to generate pseudo labels. More information on the benchmark data, baseline models, chemical representation, and evaluation metrics can be found in supplemental material.

**Figure 1.**
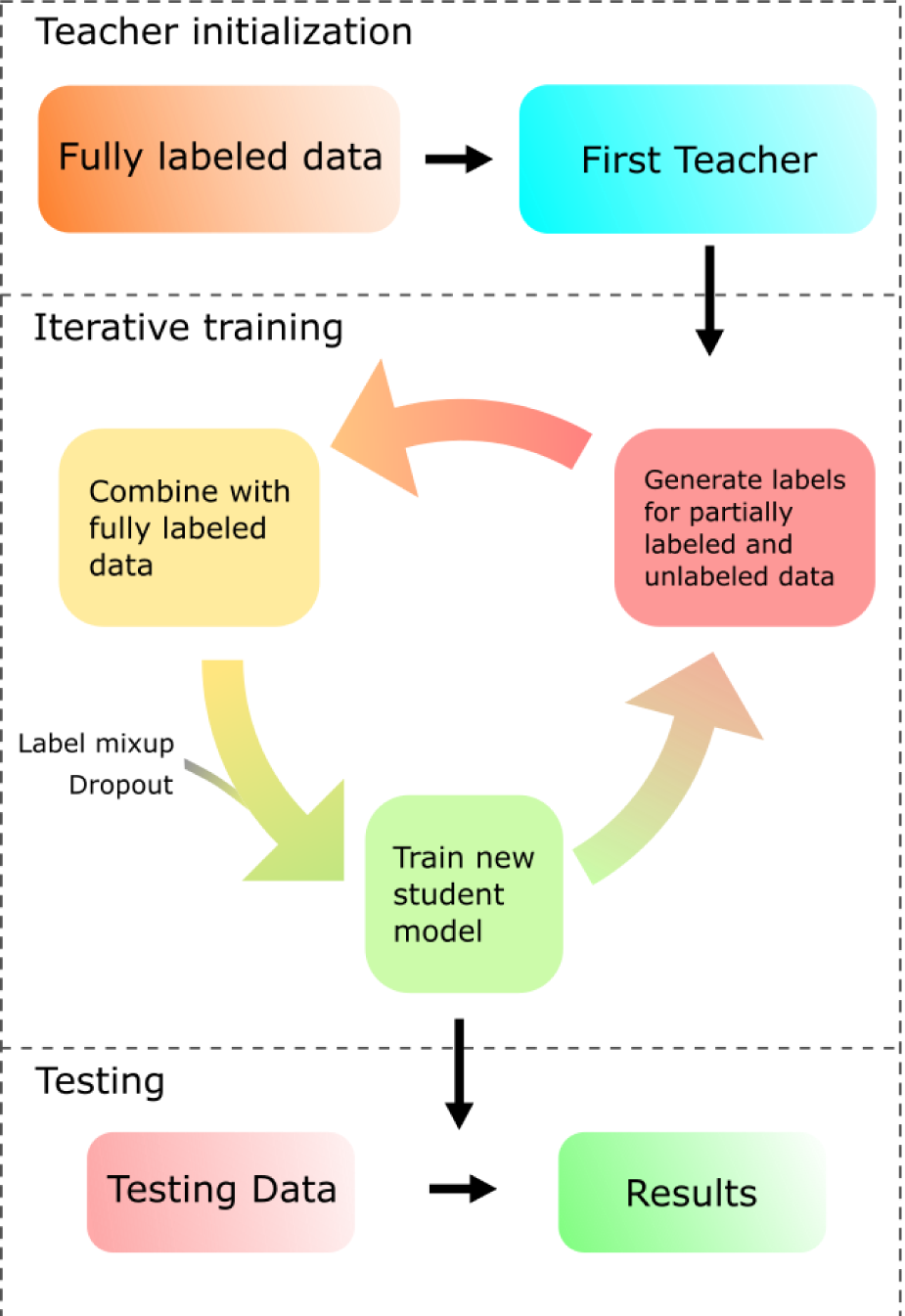
Overview of the workflow.

### 2.2. Self-training with noisy student

Knowledge distillation with self-training was used to reduce the size of models when the method was proposed (Hinton *et al*., 2015). Typical self-training uses a large teacher model trained with true/sparse labeled dataset to generate soft/continuous labels for the same dataset. One or more smaller student models are trained with the soft labeled dataset. The major reason why this model works is the continuous labels are more informative than the sparse label. They tell the student model which samples are “easier” to classify with labels that are closer to 0 or 1 and which samples are harder to classify with labels closer to 0.5. Thus, though the student models have less parameter than the teacher model, they can “memorize” similar amount of information as the teacher model and reach comparable accuracy rate. Xie et al. utilized the self-training method in an opposite way (Xie *et al*.). In their work, they used a small teacher model to train larger student models and make the student models perform not comparable but better than the teacher model. To make the method work, they introduced a large number of unlabeled images other than the labeled training images. They labeled these unlabeled outside images with the pre-trained teacher model and used these images to train the student models combined with the labeled images. In addition, another key method used in the work was introducing “noise” when training the student models by applying data augmentation to the inputs and dropout to the model parameters. In this report, we used similar strategy to recurrently train teacher and noisy student models. However, unlike images, we could not use random rotation, scaling, etc. to introduce noises into the training samples, which are chemical compounds. Thus, we used partial labeling and label mixup instead.

### 2.3. Partial labeling

To fully exploit the information in the partially labeled dataset, we use the following approach to generate pseudo labels for the partially labeled data. First, we infer the possible class of the sample based on the partial label. For example. if the label is [0, 1, _, 0, 0] where “_” represents the missing label, the possible class of the sample will only be 8 (corresponding to label [0, 1, 0, 0, 0]) or 12 (corresponding to label [0, 1, 1, 0, 0]) of the one-hot encoding. The rest digits of the one-hot label for this sample are 0. Then we predict the 32 digits one-hot label using the teacher model. Only the value at position 8 and 12 are taken from the predicted label and normalized to sum up to 1. The normalized values are filled into the missing positions to generate the integral label for the partially labeled sample.

### 2.4. Label mixup

“Noise” is a key factor to make the Noisy Student method work. Without noises, the classification accuracy drops around 0.7 percent in the ImageNet task (Xie *et al*.). However, in our case, introducing noises is not a trivial task. ECFP fingerprint (Rogers and Hahn, 2010) that is used to represent chemical (see supplemental material for details) is invariant to stereo transformations, so random rotation is not applicable. Chemicals have fixed sizes, so scaling is not applicable neither. To overcome the difficulty, we utilized an augmentation technique named mixup (Zhang *et al*., 2017). What it does is randomly pick two inputs, add both the inputs and the labels together with a coefficient sampled from beta distribution. Concretely, the mixup method can be represented by the following equations:

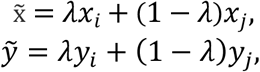

where 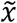 and 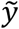 are mixed input and label. λ is a coefficient sampled from beta distribution *Beta*(α, α), α ∈ (0, +∞). *x*_*i*_ and *x*_*j*_ are the ECFP vectors of two randomly sampled chemicals. *y*_*i*_ and *y*_*j*_ are the labels of the chemicals. We tested α = 0.2 and α = 0.4 but did not find significant performance differences. For all the experiments in this report, the value of α is 0.4 if not specifically indicated.

### 2.5. Training strategy with partial labeled noisy student PLANS

We split the fully labeled CYP450s dataset into training and testing sets with a ratio of 7: With the training set, we first train a small model as the initial teacher model. Then we use the teacher model to generate pseudo labels for the partially labeled CYP450 data and unlabeled ChEMBL24 data. Followed by that, we combine fully labeled data, partially labeled data, and outside unlabeled data to train the Medium model. The trained Medium model is used to generate the labels for the partially labeled and unlabeled samples the same as described above. These regenerated samples and the fully labeled data are finally used to train the large model. Loss function used in our training is cross entropy as below:

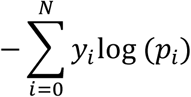

where *N* is the number of classes, *y*_*i*_ is the label of the *i*^*th*^ class, *p*_*i*_ is the predicted possibility that the sample belongs to the *i*^*th*^ class. All the training is done on a single Nvidia Tesla V100 GPU. Every experiment was repeated 5 times by random splitting the fully labeled dataset with put back. The training is terminated when the validating loss is stable. Partially labeled data and unlabeled data were only used for training.

### 2.6. Training data balancing

The labels in the CYP450s dataset are unbalanced. The number of chemicals that does not bind to any of the five CYP450s is much larger than the number of chemicals bind to at least one CYP450. In our experiment, we also tried to solve this problem by taking the advantage of self-training method. When introducing the data balancing in the experiment, the unlabeled data were not simply combined with fully labeled and partially labeled data. Instead, the procedure of adding unlabeled data to the training set was closely monitored. The sample was added to the training set only when the predicted pseudo label was not all negative and the class had less samples than the all-negative class. We remove all the unlabeled sample from training set and repeated this balancing every time when a new teacher model was trained.

## 3. Results

To assess the feasibility of the Noisy Student method in computer-aided drug design, we created a CYP450s benchmark dataset, which includes CYP1A2, CYP2C9, CYP2C19, CYP2D6, and CYP3A4 as targets, based on the experiment published in (Veith *et al*., 2009). First, we tested the prediction performance of four conventional ML algorithms and compared them to the Noisy Student trained multilayer perceptron (MLP) model. Then we introduced partially labeled noisy student (PLANS) to improve the performance. At last we balanced the training set with PLANS, and showed the balanced training set could further boost the model performance.

### 3.1. Self-training with noisy student improves the prediction of CYP450 binding profile

Other than the neural network model, we tested three types of most widely used machine learning methods, which were SVM, RF, and gradient boosting. For the gradient boosting, we chose AdaBoost and XGBoost. The configurations of the models can be found in supplementary methods session 5. Table 1 shows the classification results with the above four models.

**Table 1.** Classification results of the baseline models. The evaluation metric of the best performed model is highlighted in bold.

|  | Accuracy | Precision | Recall | F1 |
| --- | --- | --- | --- | --- |
| SVM | $54.37 \pm 1.62$ | $0.38 \pm 0.02$ | $0.84 \pm 0.01$ | $0.53 \pm 0.02$ |
| RF | $53.22 \pm 1.14$ | $0.33 \pm 0.01$ | $0.84 \pm 0.00$ | $0.47 \pm 0.01$ |
| AdaBoost | $51.34 \pm 1.17$ | $0.20 \pm 0.01$ | **$0.93 \pm 0.01$** | $0.33 \pm 0.02$ |
| XGBoost | **$54.81 \pm 1.15$** | **$0.50 \pm 0.02$** | $0.78 \pm 0.01$ | **$0.61 \pm 0.01$** |

The gradient boosting models performed better compared with SVM and RF model. It was worth noting that while the AdaBoost model got the highest recall value, its precision and F1 scores were much lower than the other three models. That means the AdaBoost model was affected the most by the data imbalance, which was a known disadvantage of AdaBoost. XGBoost, on the other hand, got well balanced precision and recall. Thus, its F1 score was the highest among the baseline models. XGBoost also showed the best accuracy.

To evaluate the feasibility of Noisy Student (NS) method, we trained an MLP model with and without NS. When using NS, the Small, Medium, and Large models as described in supplementary methods session 4 were trained iteratively with the fully labeled data and evaluation was done with the Large model. When not using NS, the Large model was trained and evaluated directly with the same training and testing sets as the model trained with NS. As shown in Table 2, all of the MLP models showed more balanced behaviors. Even the MLP without NS had got higher precision and F1 scores than the XGBoost model. When label mixup was applied, the MLP model without NS was already be able to get an almost equivalent accuracy to XGBoost while keeping a balanced performance on precision and recall. After introducing NS, the MLP models surpassed the XGBoost model with a margin of 3.0 % on accuracy and up to 13.0% on F1 score.

**Table 2.** Performance of MLP models trained without outsider data. The evaluation metric of the best performed model is highlighted in bold.

|  | Accuracy | Precision | Recall | F1 |
| --- | --- | --- | --- | --- |
| MLP | $51.97 \pm 1.12$ | $0.64 \pm 0.03$ | $0.72 \pm 0.02$ | $0.68 \pm 0.02$ |
| MLP + mixup | $54.28 \pm 0.79$ | $0.60 \pm 0.02$ | $0.73 \pm 0.01$ | $0.66 \pm 0.01$ |
| MLP + NS | $56.11 \pm 1.63$ | **$0.64 \pm 0.01$** | **$0.76 \pm 0.02$** | **$0.69 \pm 0.01$** |
| MLP + mixup + NS | **$56.48 \pm 1.45$** | $0.60 \pm 0.04$ | **$0.76 \pm 0.02$** | $0.67 \pm 0.02$ |

### 3.2. Exploitation of partial labels improves the performance of self-training with noisy student

Our CYP450s dataset contains many partially labeled samples. This is also the case for many pharmacological datasets, e.g. Tox21, ToxCast, MUV datasets in the MolNet, which is a widely used benchmark database for chemistry related machine learning projects (Ramsundar *et al*., 2019). There were studies about how to exploit partially labeled data (Yu and Zhang, 2017; Nguyen and Caruana, 2008). Generally speaking, these studies were trying to maximize the margin between candidate labels and the negative labels. NS provides a more direct way to exploit partially labeled samples. Especially when the dataset contains both fully labeled samples and partially labeled samples as our CYP450s dataset. As described in session 2.5, we utilized the teacher model trained with fully labeled data to generate semi-solid labels for the partially labeled data. Then the partially labeled data were combined with the fully labeled data to train the student model. The results were shown in Table 3.

**Table 3.** MLP trained with fully and partially labeled data. The evaluation metric of the best performed model is highlighted in bold.

|  | Accuracy | Precision | Recall | F1 |
| --- | --- | --- | --- | --- |
| MLP+PLANS | $58.94 \pm 0.96$ | $0.72 \pm 0.02$ | **$0.78 \pm 0.01$** | **$0.75 \pm 0.01$** |
| MLP+PLANS+mixup | $58.04 \pm 0.70$ | $0.69 \pm 0.02$ | $0.76 \pm 0.01$ | $0.72 \pm 0.01$ |
| MLP+PLANS+balancing | $59.02 \pm 1.12$ | **$0.73 \pm 0.03$** | $0.76 \pm 0.02$ | **$0.75 \pm 0.01$** |
| MLP+PLANS+balancing+mixup | **$59.25 \pm 1.04$** | $0.68 \pm 0.03$ | **$0.78 \pm 0.01$** | $0.73 \pm 0.01$ |

Compared with original NS-trained model, Table 3 shows that the accuracy of the PLANS-trained model improved about 5.0 % (p = 0.005) and 2.8 % (p = 0.031) without the mixup and with the mixup, respectively. Precision, recall, and F1 score also significantly increased. Especially, the precision increased 15.0 % (without the mixup) and 12.5 % (with the mixup). It is worth noting that, surprisingly, the mixup augmentation approach made the model perform worse. A possible explanation is both partial labeling and mixup introduce noises into the training set. When a partially labeled sample was added to another sample, the noise is so large that it actually confuses the model. We will leave this topic to our future work.

### 3.3. Balance the training dataset with outside data

Out training dataset is highly unbalanced. As shown in Figure 2A, the fully labeled dataset has much more all-negative data than other classes. To solve this problem, we introduced outside unlabeled dataset to balance our training set. The unlabeled data were labeled with trained teacher model and added to the training set except the samples that were labeled as all-negative. The maximum number of samples in each class was capped with the number of samples in the all-negative class. Details of data balancing can be found in session 2.6. After balancing, the labels were more evenly distributed (Figure 2B). However, the classes that had too little samples still could not be balanced to the same level as other classes. A possible solution is to introduce more noises when training the teacher models. That will be left to our future work.

**Figure 2.**
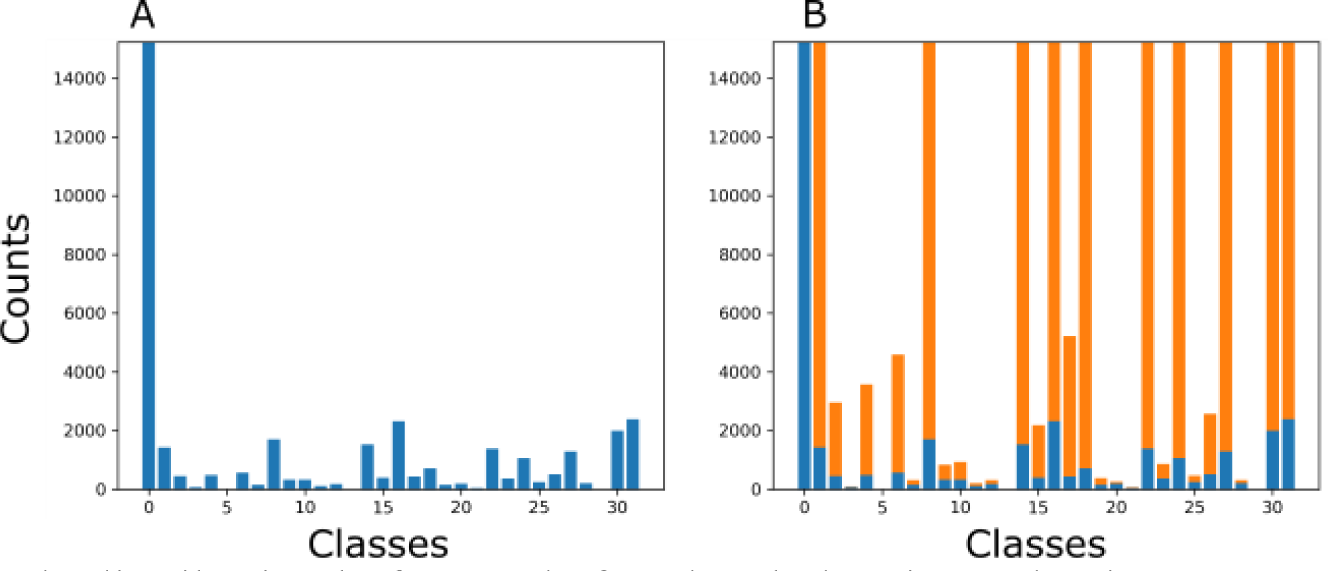
Sample distribution before and after data balancing. Blue bars represent the original samples. Orange bars represent the samples added from ChEMBL24 dataset.

The performance of the model trained with balanced training set was shown in the last two rows of Table 3. The accuracy and recall improved 2.1% (p-value=0.031) and 2.6% (p-value = 0.007), respectively, compared with the model trained with unbalanced dataset. There are no significant changes in precision and F1. The insignificant improvement by the data balancing strategy here may be due to the ignorance of label dependency. Due to the evolutionary relationships between CYP450s, the labels generated from the one-hot encoding are not evenly distributed in nature. It is subject to future studies to take the label relationship into account. In Figure 3 we analyzed the prediction results of the model trained with or without balanced data. The upper two panels showed the model trained with balanced data missed more samples that belong to the all-negative class while missed less samples in other classes. In opposite, the lower two panels showed the model made less mistakes in classifying samples into the all-negative class while making more mistakes in classifying samples into other classes. The summation of these two effects made the model perform better than the model trained with unbalanced raw data.

**Figure 3.**
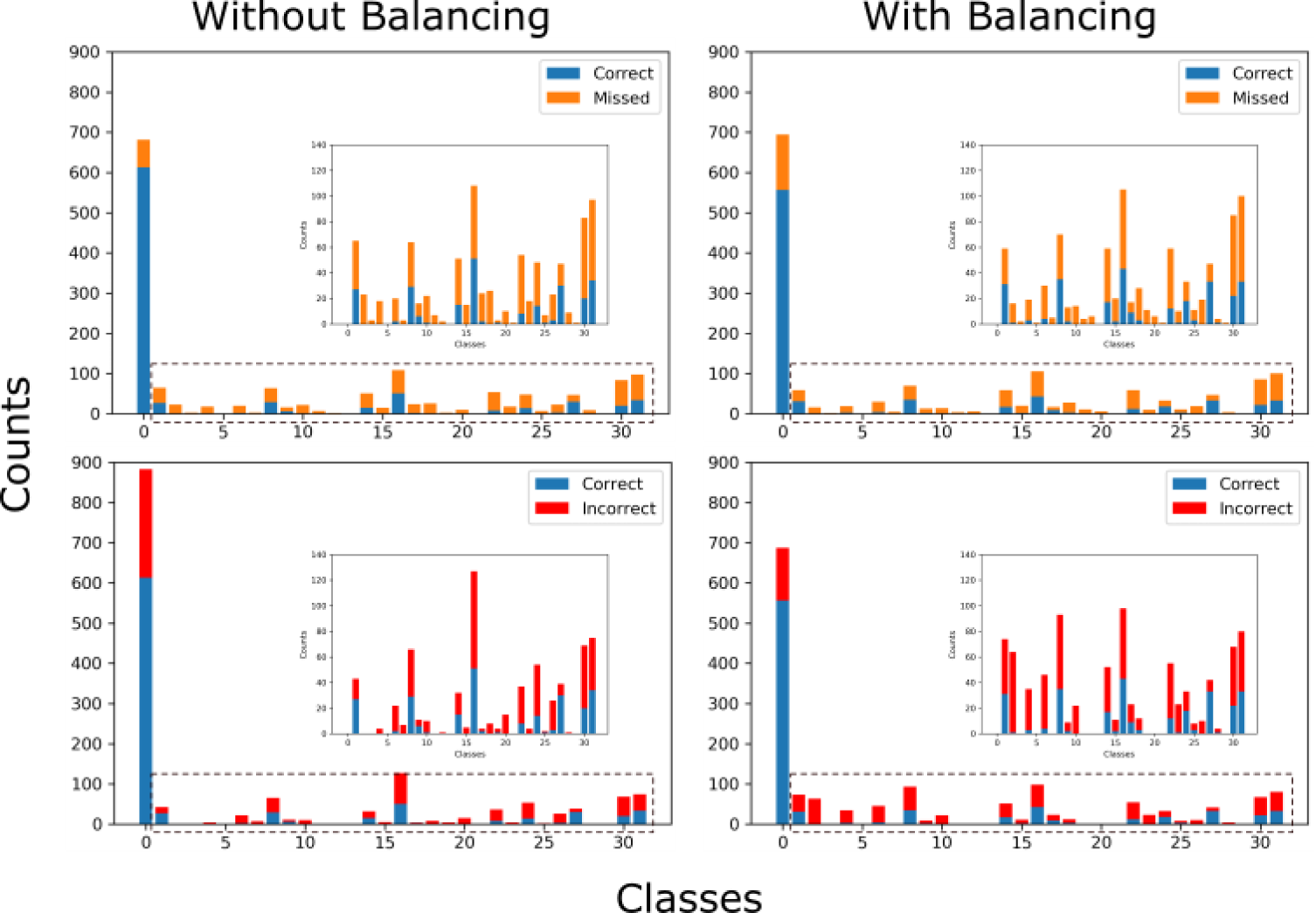
Analyzing the training results with or without the data balancing. Blue bars represent the samples that were correctly predicted. Orange bars represent the samples that the model failed to recall. Red bars represent samples that were incorrectly classified into the class by the model. The subpanels are the zoom in of the classes without the all negative class.

## 4. Discussion

Lacking of labeled data and imbalanced dataset are two general difficulties in the supervised learning with chemical and biological data. The difficulties are hard to overcome from the experiment side because of the immense cost of efforts and time for large scale experiments. That makes developing robust models that can learn useful information from small and imbalanced datasets a meaningful and important direction. In this work, we developed a self-training algorithm, Partial Labeled Noisy Student (PLANS), and applied it to an enzyme-affinity-prediction task. Our experiments proved the algorithm not only outperforms the conventional machine learning algorithms, more importantly, it also improves the performance of MLP with a big margin. Moreover, PLANS provides ideas exploiting partially labeled data as well as using unlabeled data to augment and balance labeled data. However, there is one critical problem remains unsolved.

The most important two operations of the NS method are introducing noises and utilizing abundant unlabeled data when training the student models (Xie *et al*.). However, other than random dropout, the most widely used data augmentation methods for images, like rotation and scaling, are not applicable to chemical compounds. Thus, in our work, we used mixup to do data augmentation, and method worked in most cases. Nevertheless, we noticed mixup conflicts with the usage of large amount of unlabeled data. Actually, when we used the whole ChEMBL24 dataset without capping it with the largest class in our training set together with mixup to train the model, the performance is worse than using only the mixup. The reason could be mixing model generated labels introduces more noise than useful information. We partially overcome it by only using a small portion of the unlabeled data to balance our training set. In the future, if a better solution other than mixup can be found to accomplish data augmentation task for chemical data, utilize large amount of unlabeled data should be more informative and further improve the performance of PLANS trained model.

## 5. Acknowledgements

This work is supported by National Institute of General Medical Sciences (R01GM122845) and National Institute on Aging (R01AG057555) of the National Institute of Health.

