## Supplementary Methods for "Exploration of Chemical Space with Partial Labeled Noisy Student Self-Training for Improving Deep Learning Performance: Application to Drug Metabolism"

### 1. *CYP450s Dataset*

We chose the five CYP450s described above as protein targets to create our benchmark dataset (Veith *et al.*, 2009). The Uniprot Knowledgebase ID of the five CYP450s are P08684 (CYP450 3A4), P05177 (CYP450 1A2), P10635 (CYP450 2D6), P33261 (CYP450 2C19) and P11712 (CYP450 2C9). The chemicals from the dataset were labeled with active or inactive based on their binding affinity to each CYP450 target. However, not all of the chemicals in the dataset have explicit binding affinity label to all five CYP450s. Some labels are missing because the experiments were not conducted. In this report, we call the chemicals in the CYP450s dataset with at least one missing label partially labeled data and the rest chemicals fully labeled data. The CYP450s dataset has 17121 chemicals, among which 5162 chemicals are fully labeled while 11959 chemicals are partially labeled. The full labels are converted to 32 classes labels in which each class represents a unique combination of the five affinity labels, e.g. [0, 0, 0, 0, 0] is converted to class 0, [0, 1, 1, 0, 1] is converted to class 13. To exploit the partially labeled data, we took the advantage of self-training strategy and generated partially confident labels, which will be described later. In the results session, we will show the partially labeled data improve the performance of the trained model with a big margin comparing to the model trained only with the fully labeled and the unlabeled data.

### 2. *ChEMBL24 dataset*

The ChEMBL "is a manually curated database of bioactive molecules with drug-like properties" (Mendez *et al.*, 2018). In our work, as the outside dataset for the Noisy Student method, we used the ChEMBL24, which was released in 2018 and has over 1.7 million drug-like chemicals.

### 3. *Extended Connectivity Fingerprint*

Extended connectivity fingerprint (ECFP) is a popular method for chemical structure information embedding (Rogers and Hahn, 2010). It converts chemical structures with different lengths to a fixed-length binary vector, which reserves most bonds and atom types information. Thus, ECFP is widely used as an input feature for machine learning, especially deep learning models. To minimize the frequency of bit collision, we chose a vector length of 2048, which will result in a relatively sparse vector so that the identifiers are less likely to overlap. We used RDKit to convert the simplified molecular-input line-entry system (SMILES) strings to 2048 bits vectors with the radius parameter set to 4 (Landrum, 2006)**.**

### 4. *Neural Network Model*

To prove the feasibility of Noisy Student in pharmacology field, we used the most basic MLPs to attenuate the influence from the model architectures. The Small, and also the initial teacher model that we used have seven hidden layers with 4096, 8192, 8192, 4096, 2048, 1024, and 512 hidden units, respectively. The Medium model adds two layers with 6144 hidden units on top of the Small model after layer 1 and layer 3. The Large model, in addition to the Medium model, inserts two layers with 12288 hidden units after layer 4 of the Small model. A dropout layer is added before the final output layer in all three models during the training. The rate of the dropout was kept as 0.7 in all the experiments. All the models are implemented with Tensorflow (Abadi *et al.*, 2016).

5. *Baseline models*

In addition to comparing the Large model to the Small model, we also compared the MLP model to some conventional machine learning methods, including support vector machine (SVM), random forest (RF), AdaBoost, and XGBoost. The SVM model we tested was the C-support vector machine implemented with scikit-learn (Pedregosa *et al.*, 2011). We did hyperparameter tuning and found that the SVM with regularization parameter C of 1.0 and kernel coefficient gamma as "scale" performed the best for our CYP450 task. We did the same hyperparameter tuning to the RF and AdaBoost models implemented with scikit-learn. The best RF model has max depth of 5, max features of 1024 (half the length of the ECFP), and number of estimators of 1000. The AdaBoost model used decision tree as base estimator with the number of estimators of 200 and learning rate of 0.5. The XGboost model was implemented with xgboost package (Chen and Guestrin, 2016). The XGBoost model that performed the best had learning rate of 0.1 and max depth of 14 in our experiments. All of the not mentioned parameters were left as default.

6. *Evaluation Metrics*

For our multi-class classification task, since the labels were highly unbalanced, to better evaluate our model, in addition to accuracy, we also introduced micro precision, recall, and F1 scores, which were the metrics better fit uneven dataset. To calculate these scores, the predicted one-hot labels were converted back to 5-class multi-label labels in our CYP450 task case.
